# High-risk microbial signatures are associated with severe parasitemia in controlled *Plasmodium* infections of both humans and rhesus macaques

**DOI:** 10.1101/2022.09.06.506695

**Authors:** Andrew T. Gustin, Courtney A. Broedlow, Kevin Hager, Ernesto Coronado, Solomon Wangari, Naoto Iwayama, Chul Y. Ahrens, William D. Garrison, Kathryn A. Guerriero, Kristina De Paris, Michael Gale, Nichole R. Klatt, James G. Kublin, Jennifer A. Manuzak

## Abstract

While functions of the gastrointestinal (GI) microbiome include maintenance of immune homeostasis and protection against infectious disease, its role in determining disease severity during *Plasmodium* infection has been limited to mouse models and observational human cohorts. Here, we performed controlled *Plasmodium* infection in both humans and rhesus macaques (RMs) to experimentally determine the impact of GI microbiome composition on disease progression. Through analysis of serially collected microbiome samples, we identified a high-risk microbial signature that strongly associated with increased risk of developing severe parasitemia in human participants. Importantly, we identified a parallel phenomenon in RMs. The combined weight of this evidence demonstrates that pre-infection GI microbiome composition is highly indicative of *P. falciparum* disease risk. Moreover, our observation that *P. fragile*-microbiome dynamics in RMs closely mirrors *P. falciparum*-microbiome interactions in humans strongly supports the use of this model in pre-clinical investigations of novel microbiome-targeting approaches to reduce malaria burden.

## Introduction

In 2020 there were 241 million cases of malaria worldwide, resulting in over 627,000 deaths^1^; leading *Plasmodium* infection, the causative agent of malaria, to remain a significant global public health burden. In addition to the classic clinical features of fever and anemia, *P. falciparum* malaria causes gastrointestinal (GI) manifestations, including abdominal pain, diarrhea, and vomiting^2^. Moreover, elevated GI *Plasmodium* parasite sequestration^3–6^ causes damage to GI blood vessels and contributes to disruptions in villi formation^7^, and increased epithelial permeability^7,8^, and malabsorption^9–11^. These alterations may contribute to *Plasmodium-induced* microbial translocation^8,12–15^ and dissemination of nontyphoidal salmonella (NTS)^16,17^. However, the precise biological mechanisms underlying *Plasmodium*-induced GI disruptions remain poorly defined.

GI microbes are essential in preserving epithelial barrier integrity and maintaining appropriate immune surveillance; suboptimal GI microbial consortiums influence acquisition of enteric pathogens^18^. Interestingly, GI microbiome community structure may be linked with protection against severe *Plasmodium* infection. Indeed, assessments of the fecal microbiota from Ugandan children suggest that community composition varied between children that developed severe malaria anemia versus those with asymptomatic *P. falciparum*^19^. Additionally, an observational study of children in Mali demonstrated that time to *P. falciparum* infection was significantly delayed in those with higher stool abundances of *Bifidobacterium* and *Streptococcus* species, suggesting an association between GI microbiome composition and decreased prospective risk for *Plasmodium* acquisition^20^. In mice, higher GI abundances of *Lactobacillus* and *Bifidobacterium* were associated with reduced parasite burden following *P. yoelii* challenge^21^, while decreased abundance of *Lactobacillaceae* and increased abundances of *Enterobacteriaceae* and *Verrucomicrobiaceae* were inversely and positively correlated with *P. berghei* ANKA burden, respectively^22^. Finally, differences in GI microbial communities in mice resistant or susceptible to *P. yoelii* infection were associated with variations in the quality and specificity of spleen anti-*Plasmodium* antibodies, providing a potential mechanism by which the GI microbiome could influence the severity of *Plasmodium spp*. infection^23^. In sum, these findings support the notion that the pre-existing GI microbiota composition may influence prospective risk for and severity of *Plasmodium* infection.

While observational studies have linked microbiota composition and the prospective risk for and severity of *Plasmodium* infection, there remains a critical need for controlled human malaria infection (CHMI) studies to determine the role of the microbiome in shaping *Plasmodium* infection outcome. Such studies will be important for determining whether previous cross-sectional and prospective findings are broadly applicable across age ranges. Additionally, a representative animal model is required to assess mechanistic relationships between *Plasmodium* infection and GI microbiota, intestinal inflammation, and mucosal epithelial barrier integrity. While mouse models have provided useful insights, intervention-based studies will likely require a non-human primate (NHP) model to assess the translational potential of microbiota manipulation on malaria severity. Here, we present evidence from controlled malaria infection studies in humans and NHPs that (1) confirms a strong relationship between pre-infection GI microbiota composition and the potential for severe malaria; and (2) demonstrates that NHP are an ideal animal model for basic and translational investigations of *microbiome-Plasmodium* interactions.

## Materials and Methods

### Controlled human malaria infection (CHMI) study and approval

Healthy adult participants were enrolled in a nonconfirmatory, 2-part, randomized, double-blinded, placebo-controlled, single-center study completed at the Seattle Malaria Clinical Trials Center to test the prophylactic efficacy, safety, and pharmacokinetics of KAF156, a novel imidazolopiperazine class of antimalarial drug (clinical trials registration number: NCT04072302)^24^. As reported previously, KAF156 was found to be safe, well-tolerated, and resulted in a high degree of protective efficacy from *P. falciparum* infection, which was tested through a controlled human malaria infection (CHMI) model^24^. A subset of participants that received either placebo (n=8), 20mg (n=13), or 50mg (n=14) of KAF156 post-CHMI exposure to *P. falciparum*-infected mosquitoes also provided stool samples at four distinct time points: (1) within 1-2 days of CHMI; (2) during a three-day course of treatment with the antimalarial drug Malarone (atovaquone/proguanil); (3) at the end of Malarone treatment; and (4) during CHMI and treatment follow-up (within D28 – D42). *P. falciparum* infection in study participants was monitored in whole peripheral blood using a *Plasmodium* 18S ribosomal RNA (rRNA) qRT-PCR that quantified Pf A type 18S rRNA^24–26^. Stool samples were frozen at −80°C within 24hours of collection. Upon thawing, DNA was extracted from human stool samples and used for 16s rRNA sequencing analysis, as detailed below.

### Nonhuman primate study animals, approval, and sample collection

Male Indian-origin rhesus macaques (*Macaca mulatta*; RMs; n=16) were housed and cared for at the Washington National Primate Research Center (WaNPRC) under a protocol reviewed and approved by the University of Washington Office of Animal Welfare (OWA) Institutional Animal Care and Use Committee (IACUC; Protocol 4266-13; Animal Welfare Assurance Number D16-00292). The housing, care, and procedures performed in this study were done in an AAALAC-accredited facility, in accordance with the regulations put forth by the United States Department of Agriculture, including the Animal Welfare Act (9 CRF) and the Animal Care Policy Manual, as well as with the guidelines outlined by the National Research Council in the Guide for the Care and Use of Laboratory Animals and the Weatherall Report. All RMs were provided with a commercial primate chow (Lab Diet, PMI Nutrition International, St. Louis, MO) twice per day, supplemented with daily fruits and vegetables and water ad libitum. Novel food items, foraging opportunities, and destructible and indestructible manipulanda were provided as environmental enrichment. All RMs were housed in stainless steel cages with a 12/12 light cycle. All cage pans and animal rooms were cleaned daily and sanitized at least every two weeks. Stool was collected from each RM two weeks prior to *P. fragile* inoculation (Week −2), the day of *P. fragile* inoculation (Week 0), and during *P. fragile* infection and treatment (Weeks 1, 2, 3, 4 and 6). The evening prior to stool collection, RMs in full social contact were shifted into protected (grooming) contact only. The following morning, stool was collected from each animal’s cage pan and immediately stored at −80°C. Following cage pan stool collection, RMs were returned to full social contact. Upon thawing, DNA was extracted from NHP stool samples and used for 16s rRNA sequencing analysis.

### *P. fragile* inoculation, post-infection monitoring, and treatment

*P. fragile*-infected RM erythrocytes were prepared as previously described^27^. Briefly, frozen erythrocytes from a *P. fragile*-infected RM were thawed rapidly at 37°C and transferred into a sterile tube. Erythrocytes were resuspended in 12% NaCl while gently shaking, followed by a 5min incubation at room temperature without shaking. Next, 1.6% NaCl was added dropwise while gently shaking, followed by centrifugation at 1400RPM, room temperature, 10min. After centrifugation, the supernatant was aspirated and discarded, and the pellet resuspended in a solution of 0.9% NaCl and 2% dextrose, followed by centrifugation at 1400RPM, room temperature, 10min. Finally, the supernatant was aspirated and discarded, and the pellet resuspended in 0.9% NaCl solution. RMs were intravenously inoculated with up to 20×10^6 *P. fragile*-infected erythrocytes in 1ml 0.9% NaCl solution.

Giemsa staining of thin blood smears was used to assess *P. fragile* parasitemia in each RM. Blood for the smears was collected from non-sedated RMs using positive reinforcement conditioning. Parasitemia checks occurred three times per week following *P. fragile* inoculation, with an additional check occurring at week 6 post-*P. fragile* infection. Parasitemia was quantified using a light microscope and 1:10 ratio Miller Disk Reticle (Microscope World, Carlsbad, CA), as previously described^28^. Briefly, the total number of all red blood cells (RBCs), including both infected RBCs (iRBCs) and non-infected RBCs, inside the minor square are enumerated, followed by counting of all iRBCs in the major square. A total of fifteen fields were counted in three separate areas of the slide, for a total of 45 fields. The % parasitemia was calculated in each of the three areas as follows: (# of iRBC in major square / # of all RBC in minor square) x 10. The % parasitemia in each of the three areas was then averaged together to determine the final % parasitemia for each RM at each timepoint.

The % parasitemia at each time point was used to determine the need for anti-malarial treatment. RMs were treated using the following guidelines: if parasitemia was between 0.5-1%, RMs were treated orally with quinine sulfate (150mg; NDC: 53489-0700-07; Sun Pharmaceuticals, Princeton, NJ) once per day. If parasitemia was above 1%, RMs were treated orally with quinine sulfate twice per day. If parasitemia was above 0.5% for two consecutive weeks despite quinine sulfate treatment, RMs were switched to oral chloroquine treatment (10-20mg/kg; NDC: 64980-0178-02; Rising Pharmaceuticals, East Brunswick, NJ) once per day. Quinine sulfate and chloroquine treatments were halted as soon as RMs were observed to be below the 0.5% parasitemia threshold.

### 16s rRNA sequencing

DNA was extracted from both NHP and human stool samples using the PowerFecal DNA Isolation Kit (Qiagen, Valencia, CA). Sequencing of the 16s small subunit ribosomal ribonucleic acid (SSU rRNA) gene was performed at the University of Minnesota Genomics Center. Briefly, a quality control qPCR was performed in triplicate to verify that sufficient quantities of the 16s SSU rRNA gene were present in the sample for sequencing. After confirmation that adequate levels of the 16s SSU rRNA gene were extracted, the following primers were used to sequence the V3/V4 region of the 16s SSU rRNA gene: 357F (5’-CCTACGGGNGGCAGCAG-3’) – 806R (5’-GGACTACNVGGGTWTCTAAT-3’)^29–32^. Next, a dual-indexing qPCR was used to barcode and add indexes for gene amplicon sequencing^33^. The resulting amplicons were normalized, pooled, cleaned, and quantified (KAPA quantification, KAPA Biosystems, Wilmington, MA). An Illumina MiSeq (Illumina, San Diego, CA) was used to sequence the library with a 2×300 bp cycle with 15% PhiX (Illumina). All raw sequence data have been deposited in the NCBI SRA (accession number pending).

### Bioinformatics and statistical analysis

16S rRNA amplicon sequencing reads were demultiplexed in Illumina Basespace. Fastq files were assessed for read quality, trimmed (trimLeft=12, truncLen=150), dereplicated, and filtered using the dada2 pipeline in R studio. Following removal of chimeric sequences, reads were assigned a taxonomy using greengenes (gg_13_8_train_set_97.fa.gz) and a phylogenetic tree was built using MSA and Phangorn. The reads, metadata, and tree were then combined into a phyloseq object for additional quality controls and filtering, which included removing ASVs mapping to mitochondria, chloroplast and cyanobacteria reads, as well as any ASV detected in only one sample. The resulting phyloseq object was used for all downstream analyses.

Unsupervised hierarchical clustering was conducted on center log ratio (CLR)-transformed read counts, using a divisive approach (Cluster::diana). The same CLR-transformed read counts were used to visualize human reads in PCA space. Alpha diversity was calculated using the microbiome package to assess raw reads. Prevalence, proportion tests, and CLR-transformed abundances were assessed using custom scripts that are available upon request. Differential abundance was performed using DESeq2 to assess raw reads on genera-level objects generated using phyloseq::tax_glom. Time course data were smoothed using default options for ggplot2::geom_smooth. PCoA of RM samples were generated by first running phyloseq::ordinate on the phyloseq object, and the resulting objects were visualized using phyloseq::plot_ordination. All code is available upon request. Log2 peak parasite burdens, Shannon diversity levels, and CLR-transformed counts were compared using a pairwise Wilcoxon test. Significant differences in prevalence were assessed by proportion test. Differential abundances were determined using DESeq2 with a Wald test.

## Results

### Pre-infection human microbiome composition is associated with parasite burden during controlled malaria infection

We first assessed the stool microbiota in healthy adults participating in a CHMI study^24^. As previously reported, study participants were inoculated via infected mosquito bite (n=35). Participants that progressed to detectable parasitemia by qPCR (n=32) were included in the sequencing analysis performed here^24^. Malarone (atovaquone/proguanil) treatment was initiated according to predetermined criteria of two consecutive positive qRT-PCR test results, with at least one measurement greater than 250 parasites/ml^24^. Stool samples were collected (1) prior to challenge, (2) at peak parasitemia (prior to antimalarial treatment thresholds), (3) post-treatment, and (4) during follow-up (Fig. 1A).

**Figure 1.**
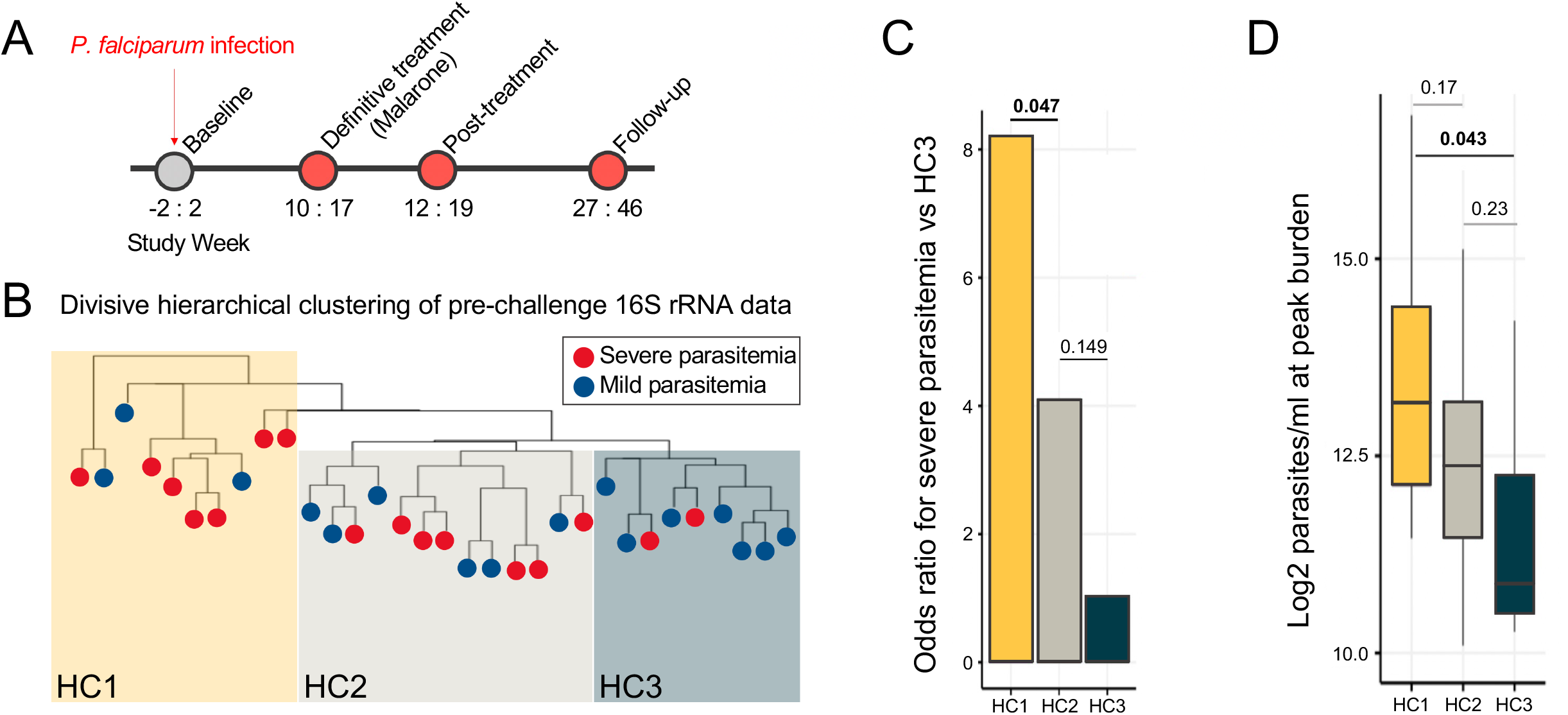
Unsupervised hierarchical clustering of 16s rRNA sequences identifies association between GI microbiota composition and risk of severe *P. falciparum* parasitemia. (A) Representative timeline depicting CHMI sample collection schedule. CHMI participants provided stool samples near *P. falciparum* challenge (between days −2 – 2), at peak parasitemia near when treatment was initiated (days 10 – 17), post-treatment (days 12 – 19), and at a follow-up visit (days 27 – 46). (B) Divisive hierarchical clustering to identify major clusters in baseline microbiota samples. Major clusters were defined as human cluster 1 (HC1; gold), human cluster 2 (HC2; gray) and human cluster 3 (HC3; dark cyan). Branch tips indicate the parasitemia severity of each participant, which was based on detection of peak parasite levels. (C) Odds ratio assessment to determine likelihood of participants within each microbiota cluster progressing to severe parasitemia. (D) Log2-transformed parasitemia values for participants in each microbiota cluster.

We hypothesized that the magnitude of *Plasmodium* parasite burden would vary according to microbiota composition. To test this, we performed unsupervised, divisive hierarchical clustering of 16S rRNA gene sequencing data to identify major community types. Baseline microbiota compositions could be segregated into three distinct clusters (Fig.1B). We next categorized infection outcome as either mild or severe parasitemia, defined as parasite burden above or below the median parasitemia value. Participants in human microbiome cluster 1 (HC1; gold) were more than eight times as likely to experience high parasitemia as HC3 participants (dark cyan; odds ratio (OR) = 8.17; p = 0.047; 95% confidence interval (CI) = 1.03 – 64.94), while HC2 participants (gray) were more than four times as likely to develop high parasitemia than HC3 participants (OR = 4.08; p = 0.149; 95% CI = 0.60 – 27.65) (Fig. 1C). When assessing parasite burden in each cluster, HC1 participants developed significantly higher levels of parasitemia as compared to HC3 participants (p=0.043); HC1 peak parasitemia was also elevated compared to HC2, although this difference did not reach statistical significance (p=0.17; Fig. 1D). Parasite burden was non-significantly elevated in HC2 compared to HC3 participants (p=0.23; Fig. 1D). Importantly, we observed no relationship between microbiome composition and post-exposure treatment with KAF156 or placebo following *P. falciparum* challenge (Fig. S1A and B). Taken together, microbiota composition prior to *P. falciparum* exposure was significantly associated with parasite burden in humans.

### Human GI microbiome composition is robustly associated with malaria severity and progression

We next identified structural and compositional distinctions between microbiota clusters by performing components analysis (PCA) on all collected timepoints. HC1 participants were widely dispersed, HC2 participants were modestly dispersed, and HC3 participants were tightly clustered in PCA space (Fig. 2A and B), indicating greater variability in microbial composition in HC1 than HC2 or HC3. These patterns were especially evident with information from the third principal component added in 3-dimensional space (Fig. S2A and B). When considering baseline only, HC2 and HC3 had significant overlap along PC1 and minimal spread along PC2, while HC1 demonstrated higher variability along both PC axes (Fig. 2C and D). These results affirmed the hierarchical cluster assignments by illustrating the degree to which HC1 differed from HC2 and HC3 in composition and variability at baseline and throughout *P. falciparum* infection.

**Figure 2.**
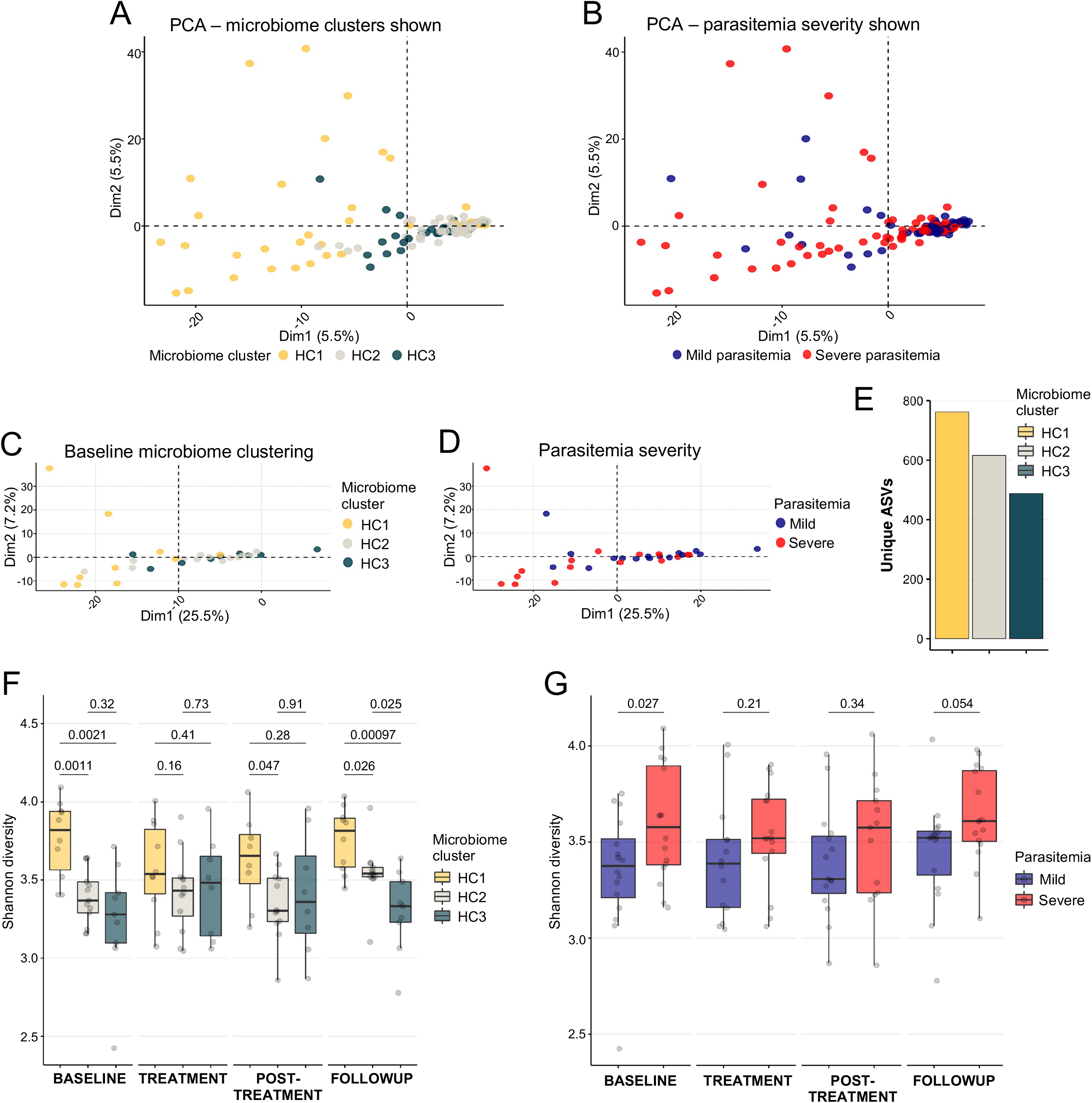
High diversity GI microbiota is linked to likelihood of developing severe *P. falciparum* parasitemia. (A) Principal component analysis (PCA) of CLR-transformed count data. Colors indicate hierarchically determined microbiota clusters, with HC1 shown in gold, HC2 in gray, and HC3 in dark cyan. (B) Replicate PCA as shown in A, with colors identifying samples associated with parasite levels. Blue points indicate mild parasitemia, while red points indicate severe parasitemia. (C) PCA of samples at baseline only, with points representative of HC1 (gold), HC2 (gray), and HC3 (dark cyan). (D) Replicate PCA of samples at baseline only with points shaded to represent severe (red) and mild (blue) parasitemia. (E) Total number of unique ASVs observed within each microbiota cluster. (F) Shannon diversity scores for participant samples at all time points, grouped by microbiota cluster. (G) Shannon diversity scores for participants at all time points, grouped by parasitemia severity.

To determine whether within-sample alpha diversity was a distinguishing aspect of cluster identities, we assessed the cumulative number of unique amplicon sequence variants (ASVs) detected at any time in each microbiota cluster. The number of unique ASVs in HC1 (n=762) was almost two-fold greater than in HC3 (n=488; Fig. 2E), indicating a greater range of taxa and explaining the wide dispersion of HC1 samples in PCA space. Additionally, HC1 Shannon diversity was elevated as compared to HC2, a finding that remained stable across all measured time points and was statistically significant at baseline, post-treatment, and follow-up (p=0.0011, 0.047, and 0.026, respectively; Fig. 2F). Likewise, measured Shannon diversity in HC1 was significantly higher than HC3 at baseline and follow-up time points (p=0.0021 and 0.00097, respectively; Fig. 2F). Shannon diversity scores for HC2 and HC3 participants were similar over time, except follow-up where HC2 showed significantly higher scores than HC3 (p=0.025; Fig. 2F). Participants with high levels of parasitemia had consistently higher Shannon diversity as compared to participants with mild parasitemia (Fig. 2G). This difference was statistically significant pre-challenge (p=0.027) and trended towards significance at follow-up (p=0.054; Fig. 2G). Taken together, elevated GI microbiota diversity prior to *P. falciparum* challenge could contribute to the development of high parasitemia during infection. Additionally, stable diversity over time suggests that pre-existing community structure is reestablished following transient changes induced by *Plasmodium* infection.

### Genus-level distinctions in human intestinal microbial composition are linked with malaria disease severity and progression

To test whether the increased microbial richness in HC1 participants would result in reduced detection and measured abundance of core ASCs compared to HC3 participants, we measured the prevalence of all ASVs detected in at least one third of all participant samples at all time points and performed proportionality tests. Highly prevalent ASVs, defined as ASVs detected in more than 75% of all samples, were consistently detected at lower frequencies in HC1 than HC3 (Fig. 3A). Conversely, ASVs with lower overall prevalence were more likely to be detected in HC1 than HC3. Among differentially prevalent ASVs, several key GI microbiota were significantly less prevalent in HC1 participants, including *Bifidobacterium longum* (p<0.0005), *Bifidobacterium adolescentis* (p<0.005), and *Akkermansia muciniphila* (p<0.005; Fig. 3A). We further assessed whether the abundances of core taxa, when detected, were different between participant groups by comparing center log ratio (CLR)-transformed count data for the 20 most prevalent taxa. Highly prevalent ASVs were significantly less abundant in HC1 participants as compared to HC2 and HC3 (Fig. 3B). The application of strict Bonferroni corrections demonstrated the strong significance of this observation and highlighted reduced levels of *Bifidobacterium adolescentis* (p=0.00069; Fig. 3B). Taken together, core GI microbiota ASVs are less prevalent and abundant in HC1 participants with high parasite burdens, and their absence is accompanied by increased detection of low prevalence taxa. These findings were robust even when accounting for decreased taxa prevalence.

**Figure 3.**
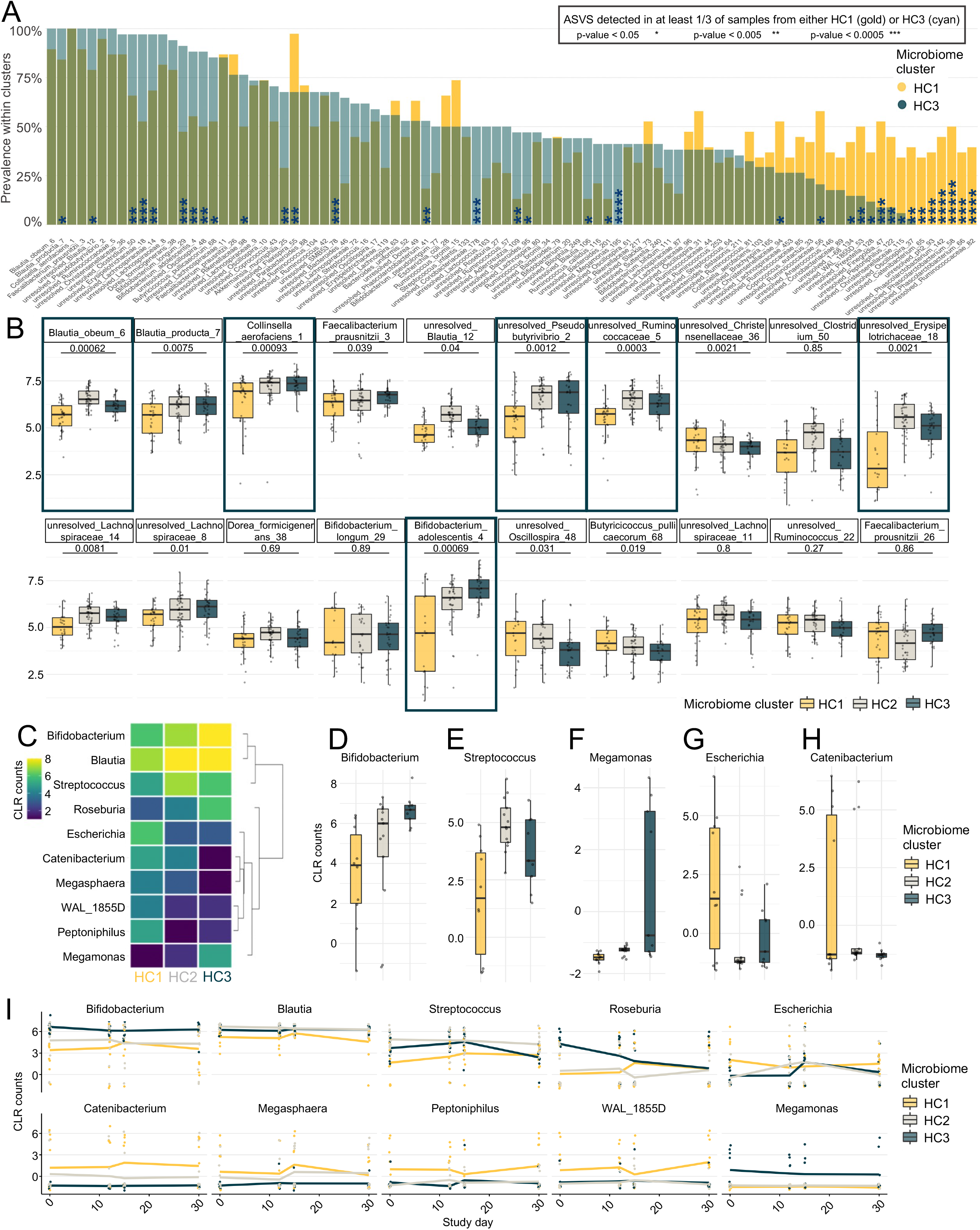
Abundance of core taxa is reduced in participants likely to develop severe *P. falciparum* parasitemia. (A) Prevalence of all ASVs that were detected in at least one third of samples from each microbiota cluster. Statistical values indicate significant differences in prevalence as determined by a proportionality test. * = p<0.05, **=p<0.005, ***=p<0.0005. (B) CLR-transformed abundance data for top 20 most prevalent ASVs, when detected. Statistical significance between HC1 and HC3 was conducted using a Wilcoxon test, with results that remained significant following Bonferroni correction boxed in dark cyan. (C) Heatmap of CLR-transformed count data for all genera detected as differentially abundant at the baseline time point, with hierarchical clustering of genera shown to the right. (D-H) CLR-transformed counts for all baseline samples in select differentially abundant genera. (I) Smoothed abundance data over time. Solid colored lines represent mean count levels. For panels A-B and D-I, HC1 is shown in gold, HC2 is shown in gray and HC3 is shown in dark cyan.

To determine whether the risk for progression to severe malaria was driven by high-level community composition differences, we performed differential abundance analysis on all pre-challenge samples at the genus level. HC1 participants had reduced baseline levels of *Bifidobacterium*, *Streptococcus*, and *Megamonas* relative to HC2 and HC3 participants (0.0067, 0.01, 0.0037, respectively; Fig 3C–F). These reductions were balanced by increased levels of *Escherichia* and *Catenibacterium* (p=0.0016 and <0.00001, respectively; Fig. 3C and G–H), among others. These baseline differences in community composition were consistent throughout infection, with levels of *Megamonas* and *Bifidobacterium* stably elevated in HC3, while *Blautia* and *Streptococcus* remained consistently highest in HC2, with HC3 approaching similar levels (Fig. 3I). Conversely, *Catenibacterium, Megasphaera, Peptoniphilius*, and *WAL_1855D* were consistently elevated in HC1 (Fig. 3I), indicating a potential association with severe malaria risk. *Escherichia* and *Roseburia* were disrupted following challenge, and although *Escherichia* returned to pre-challenge levels in all community clusters*, Roseburia* was consistently diminished in HC3 following infection. Thus, high-level distinctions between microbiota clusters contributed to risk for malaria severity, and GI microbial communities were generally stable subsequent to *P. falciparum* infection.

### Distinct RM microbial cluster is significantly associated with parasite burden during *P. fragile* infection

To further test our hypothesis that the GI microbiome plays a direct role in *Plasmodium* progression, we conducted experimental infection of rhesus macaques (RMs) with *P. fragile*, a macaque *Plasmodium* previously shown to induce clinical signs in RMs comparable to those caused by *P. falciparum* in humans^34–36^. Stool microbial communities of male RMs (n=16) were characterized two weeks prior to *P. fragile* challenge and through six weeks post-infection (Fig. 4A). Unsupervised hierarchical clustering revealed two major microbial community branches, macaque cluster 1 (MaC1) and 2 (MaC2). MaC1 RMs were 21 times as likely to develop high parasite loads as MaC2 RMs (OR = 21.00; p = 0.024; 95%CI = 1.50-293.271). Moreover, MaC1 RMs reached peak parasite loads nearly three times greater than MaC2 RMs (p=0.003; Fig. 4C). Thus, the pre-infection microbiota composition of RMs was associated with the likelihood of developing severe malaria, similarly to human CHMI participants.

**Figure 4.**
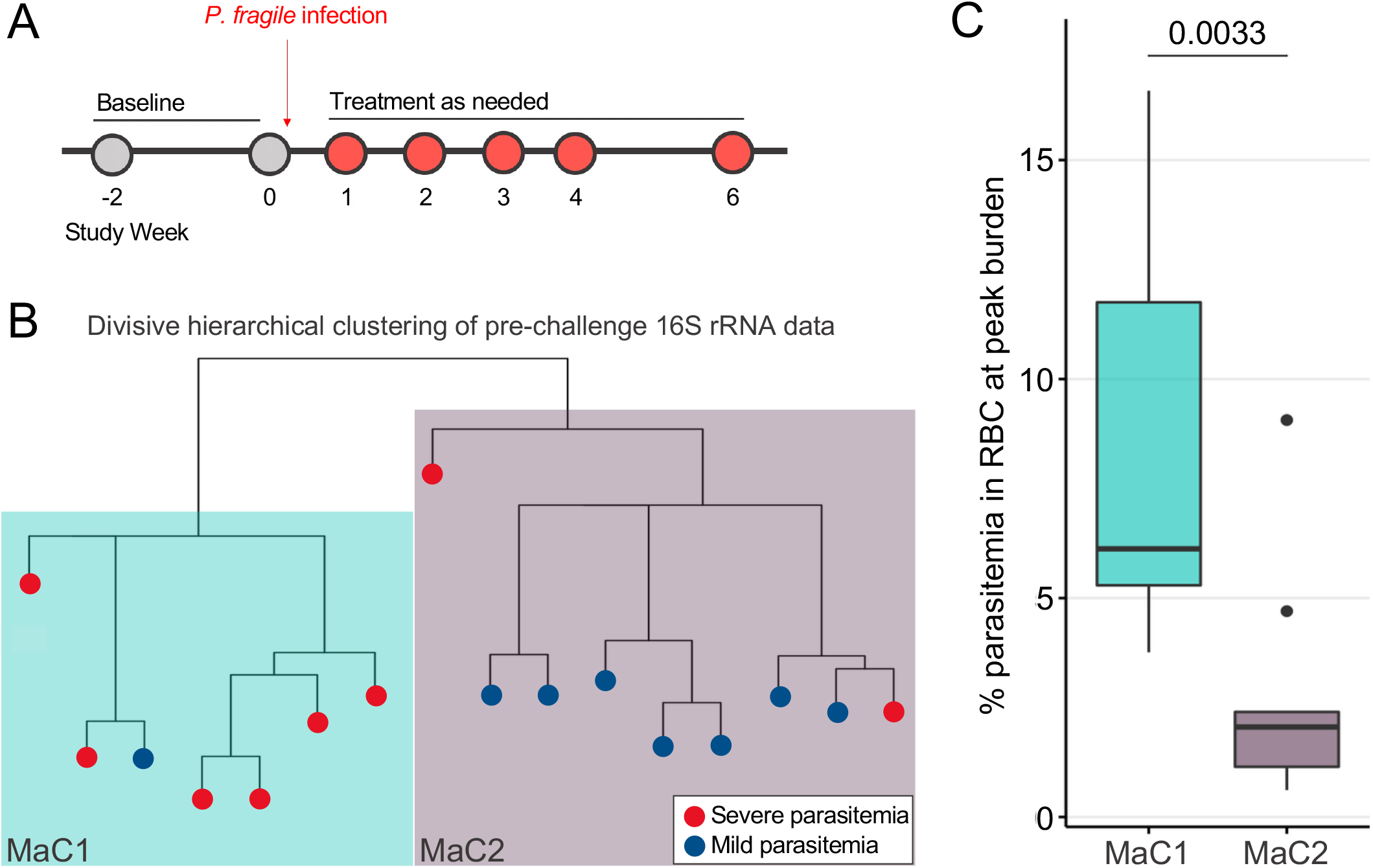
Unsupervised hierarchical clustering of 16s rRNA sequences identifies association between GI microbiota composition and risk of severe *P. fragile* parasitemia. (A) Representative timeline depicting RM sample collection schedule. RM stool samples were collected twice prior to *P. fragile* challenge, then each of the next four weeks, with final follow-up at six weeks post-challenge. (B) Divisive hierarchical clustering to identify major clusters in baseline microbiota samples. Major clusters were identified as macaque cluster 1 (MaC1; teal) and macaque cluster 2 (MaC2; purple). Branch tips indicate the parasitemia severity of each macaque, which was based on detection of peak parasite levels. (C) Log2-transformed parasitemia values for macaques in each microbiota cluster.

### RMs with distinct intestinal microbial composition have greater risk of developing severe malaria

As with CHMI data, PCA analyses reiterated the clear linkage between microbiota composition and malaria progression (Figs. S3A and B). However, unlike CHMI participants, these distinctions were not explained by alpha diversity differences. Indeed, Shannon diversity measurements indicated no significant differences between MaC1 and MaC2 at any timepoint, nor were differences observed when assessing Shannon diversity according to parasite burden (Fig. S3C and D), suggesting that distinctions between RM microbiota clusters may not be driven by differences in community organization. We next assessed community composition using PCoA of weighted-UniFrac values and found that MaC1 clustered separately from MaC2 (Fig. 5A) and displayed the clear association between microbiota composition and parasite burden (Fig. 5B). Faceting data according to timepoint demonstrated that pre-challenge microbial profiles remained distinct following recovery from *P. fragile* infection (Fig. 5C). Notably, despite this overall stability, samples from week 3 post-challenge were not distinct in PCoA space, suggesting that while microbial communities may be relatively resistant to influence from *P. fragile* infection, they may be transiently disrupted (Fig. 5C). Taken together, our data demonstrate that parasite burden was primarily driven by major compositional differences, rather than variable levels of microbiota found in both groups.

**Figure 5.**
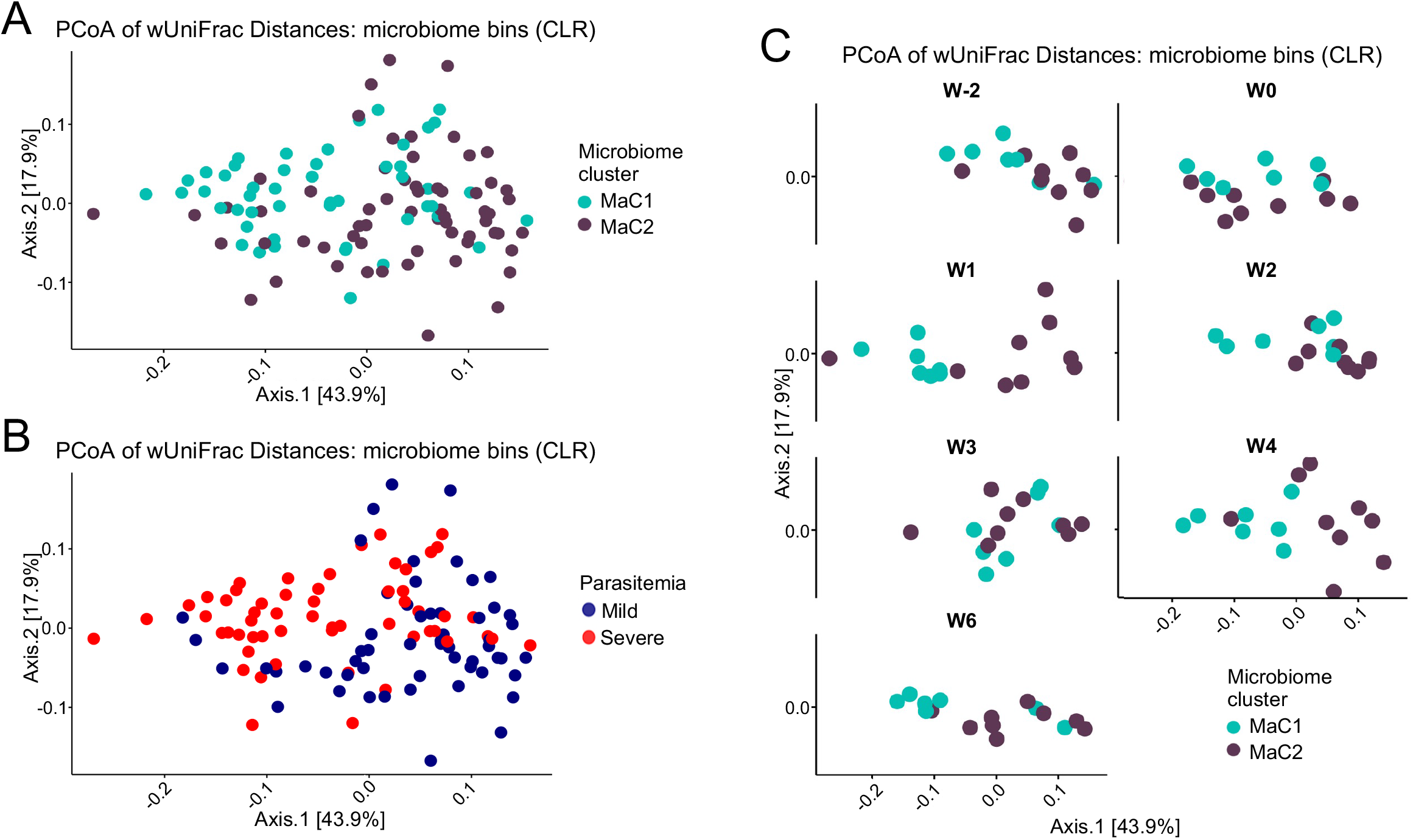
GI microbiota composition linked to likelihood of developing severe *P. fragile* parasitemia. (A) Principal Coordinate Analysis (PCoA) weighted-unifrac distances between samples. Colors indicate hierarchically determined microbiota clusters, with MaC1 shown in teal and MaC2 shown in purple. (B) Replicate PCoA as shown in (A), with colors identifying samples associated with parasite levels. Blue points indicate mild parasitemia, while red points indicate severe parasitemia. (C) PCoA of samples faceted by timepoint and colored by microbiota cluster.

### Genus-level distinctions in RM intestinal microbial composition are linked with malaria disease severity and progression

We next measured the abundance of core ASVs present in at least one third of all samples for each microbiota cluster and performed proportionality tests (Fig. 6A). ASVs from the *Prevotella* genus and *Ruminococcaea* family were more likely to be present in MaC2 RMs (Fig. 6A). Conversely, MaC1 RMs more frequently carried ASVs from the *Lachnospiraceae* family, and the *Lactobacillus* and *Treponema* genera (Fig. 6A). Notably, the majority of core ASVs had abundances that were relatively similar between MaC1 and MaC2. Two important exceptions were the elevated abundance of several *Prevotella* ASVs in MaC2 RMs, which were countered by significant elevations of *Lactobacillus* ASVs in MaC1 RMs (Fig. 6B). When assessing cumulative differences in ASVs at baseline, we identified 16 genera whose abundances differed significantly between MaC1 and MaC2 (Fig. 6C). Interestingly, the *Prevotella* genus was significantly elevated prior to challenge in MaC2 RMs (Fig. 6D), whereas MaC1 RMs had elevated baseline levels of *Ruminococcus, Treponema*, and *Sphaerochaeta* (Fig. 6E–G). Finally, pre-challenge elevations in *Prevotella* abundance in MaC2 were maintained over time (Fig. 6H), suggesting a beneficial role of the *Prevotella* genus in the context of *Plasmodium* infection, as reduced levels were consistently associated with the risk of severe parasitemia. Other genera shown to be significantly different pre-challenge varied widely in abundance throughout *P. fragile* infection, with ratios between MaC1 and Mac2 frequently inverted during weeks 1 - 3 post-infection (Fig. 6B), the period during which parasite levels were highest (Fig. S4). This observation agrees with our week-by-week breakdown of UniFrac distances (Fig. 5C), indicating that *P. fragile* infection may disrupt the GI microbiome in RMs. Notably, the abundance of most genera disrupted following *P. fragile* infection returned to pre-infection levels by 6 weeks post-challenge, thus reestablishing the distinction between MaC1 and MaC2 clusters. Taken together, similarly to humans, RMs possess natural variation in GI microbiota that appears to predispose a portion of the population to more severe *P. fragile* infection.

**Figure 6.**
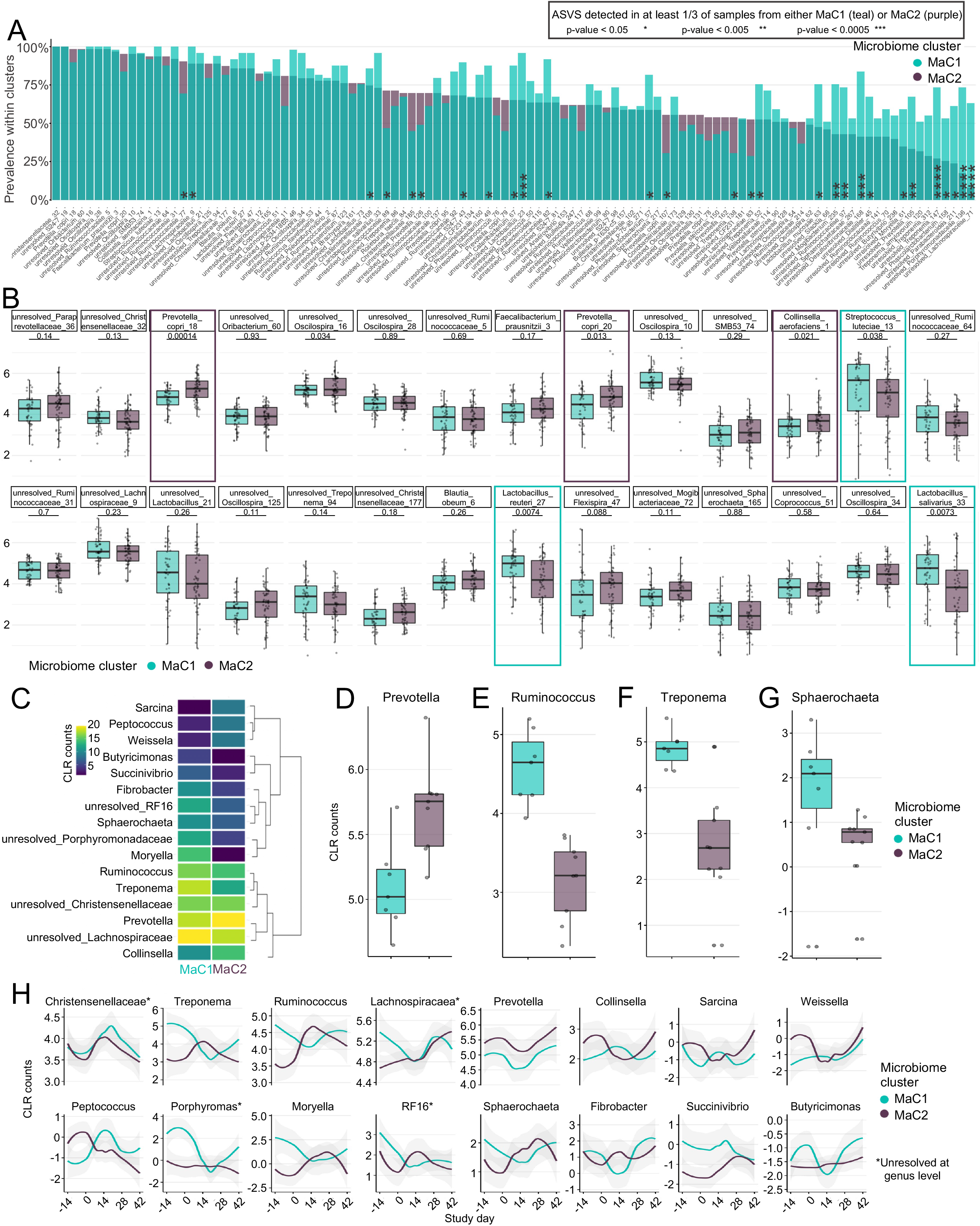
Abundance of core taxa reduced in macaques likely to develop severe *P. fragile* parasitemia. (A) Prevalence of all ASVs that were detected in at least one third of samples from each microbiota cluster. Statistical values indicate significant differences in prevalence as determined by a proportionality test. * = p<0.05, **=p<0.005, ***=p<0.0005. (B) CLR-transformed abundance data for top 28 most prevalent ASVs, when detected. Statistical significance between MaC1 and MaC2 was conducted using a Wilcoxon test, with results that remained significant following Bonferroni correction boxed in teal when abundances were significantly greater in MaC1 and purple when abundances were significantly greater in MaC2. (C) Heatmap of CLR-transformed count data for all genera detected as differentially abundant at the baseline time point, with hierarchical clustering of genera shown to the right. (D-E) CLR-transformed counts for all baseline samples in select differentially abundant genera. (F) Smoothed abundance data over time. Solid colored lines represent mean count levels. For panels A-B and D-F, MaC1 is shown in teal, MaC2 is shown in purple.

## Discussion

Here, we observed that baseline GI microbiota composition in humans was strongly associated with the potential for severe *P. falciparum* infection. Our findings are in agreement with observational studies using incident detection in human children in malaria endemic areas^19,20^ and previous murine studies^21–23^, both of which have suggested a link between GI microbiome composition and risk for severe malaria infection. Importantly, our work extends these observations to adult humans and highlights *Bifidobacterium* as a defining microbial feature in limiting risk for severe *P. falciparum* parasitemia. The agreement between studies is impressive when considering that our CHMI study was performed on a malaria-naïve cohort of adult individuals in Seattle, Washington, whereas the previous human studies were conducted among infants and children in West Africa, where malaria incidence is seasonal. These differences highlight the robust nature of our findings and position *Bifidobacterium* as a key taxon for future studies aimed at mitigating malaria severity in endemic regions.

Our data suggest that elevated diversity in the GI microbiota prior to *P. falciparum* exposure could contribute to the development of high parasitemia. Mechanistically, the loss or reduction of key commensals including *Bifidobacterium* and *Akkermansia* could result in inappropriate or inadequate priming of host immunity due to disruptions in beneficial, low-level immune stimulation^37–40^. Alternatively, their displacement by an expansion of uncommon taxa could disturb GI metabolic networks and subsequently promote pathologic changes to the GI barrier^41^. In either hypothetical, host immune disruptions could manifest locally and systemically such that *Plasmodium* replication is promoted. Recent work has also demonstrated a propensity for certain bacterial species to leverage within host evolution to promote their translocation to the liver, altering immune homeostasis in this tissue^42^. Future investigations focusing on the translocation potential of the differentially abundant taxa identified in this study are needed. Alternatively, microbial composition may serve as a proxy for indirectly related facets of overall health that may ultimately play larger roles in malaria progression; our identification of a similar phenomenon in healthy RMs, however, argues against this. Finally, although unlikely, microbiome constituents may directly impact the *Plasmodium* lifecycle. Future investigations to determine the precise mechanisms by which GI microbiota composition influences the severity of *P. falciparum* infection in humans are warranted.

As in humans, GI microbiota composition in RMs was significantly associated with severe *P. fragile* parasitemia risk. In particular, pre-challenge differences in *Prevotella* between MaC1 and MaC2 were maintained over time, suggesting that ASVs within this genus have a stable niche space that is diminished in RMs likely to develop severe parasitemia. Notably, although the GI microbiome was generally resilient to transient changes during *Plasmodium* infection in both humans and RMs, our data showed that when disruptions occurred, they were observed near peak parasitemia. These findings are consistent with previous work, which revealed only minimal differences in stool community structure and composition in Kenyan infants before and after *P. falciparum* infection and treatment^43^. However, our data are in opposition to murine studies which showed that *P. yoelli* infection resulted in decreased abundance of Firmicutes and increased abundance of Bacteroidetes^5,44^, while *P. berghei* ANKA infection resulted in decreased Bacteroidetes and Verrucomicrobia phyla, but substantially increased Firmicutes and Proteobacteria^45,46^. Taken together, the striking degree to which our RM GI microbiome findings agree with both our CHMI data and prior human observational studies supports our hypothesis that microbiome composition is a significant determining factor in *Plasmodium* infection. Moreover, our findings indicate the significant translational potential of the RM *P. fragile* model. Given the additional strong similarities between RMs and humans in terms of GI functionality and mucosal immunity, future studies utilizing this model to investigate the impact of *P. fragile*-mediated GI microbiome disruptions on intestinal mucosal barrier integrity and malaria disease progression are justified.

A major strength of our RM model is that it allowed for a greater degree of control over critical experimental and population variables. Indeed, although our CHMI study employed narrowly defined parameters for recruitment^24^, there remained a potential for unassessed genetic, nutritional, and social factors which may have contributed to microbiota variability. This created a challenge for determining whether observed differences in baseline GI microbiota composition were a proxy for some other aspect of *P. falciparum* susceptibility, or alternatively, played a direct role in risk for more severe infection. The RM model allowed us to overcome some of these challenges, including the reduction of social, economic, and medical confounders common among human cohorts. Additionally, while diet is a major contributor to GI microbiota composition in both humans and RMs^47^, the RMs included in this study were fed a standardized diet of commercial primate chow and daily fruits and vegetables. Thus, while animal-to-animal differences in GI microbiota composition prior to *P. fragile* challenge were expected, this variability was unlikely to indicate unappreciated comorbidities or other factors that may have influenced *P. fragile* infection. Moreover, the ability to stratify RMs according to pre-infection microbiota composition without the added complications of needing to account for diet or other population-specific confounders, strengthens our observation of a significant role of the GI microbiome in protection against severe *Plasmodium* infection and verifies the value of this critical model system for the examination of *Plasmodium*-microbiome interactions.

A critical aspect of our study was the use of an analytical approach that leveraged divisive hierarchal clustering, as opposed to agglomerative, which enhanced our ability to segregate microbiota communities into meaningful clusters. The ability to then identify stepwise increases in peak parasite burdens across these microbiota clusters allowed us to verify previous studies suggesting an association between the microbiome and the magnitude of *Plasmodium* infection, and extend these findings to a human adult cohort, as well as a physiologically relevant RM model system. In sum, the data presented here and our specific bioinformatic approach confirms the generalizability of our observations beyond specific participant populations, as well as to other host and *Plasmodium* species.

In summary, the data presented here demonstrates that preexisting GI microbiome composition is highly indicative of *Plasmodium* disease progression in humans and RMs. Our findings decisively support the existence of a broad link between GI microbiome communities and *Plasmodium* infection severity and represent a critical extension of previous murine and human studies^19–23^ into adult human populations. Our ability to recapitulate this phenomenon in a relevant RM model expands the ability to perform mechanistic investigations of microbiota-influenced anti-malarial immunity. While our current study is limited in its ability to make mechanistic determinations on how the GI microbiome contributes to protection against severe malaria, the identification of high-level taxonomic differences in both human and RM cohorts suggests that microbiome targeting therapies, such as microbiome modulation or small molecule supplementation, could be investigated to mitigate the severity of *Plasmodium* infection. Taken together, our data provide a strong foundation for future RM and human studies to identify the functional role of distinct microbial communities in controlling malaria severity, and pre-clinically test preventative and interventional strategies leveraging the microbiome to reduce malaria burden.

## Acknowledgements

We thank the individuals that participated in the KAF156 CHMI study. We thank all veterinary and research support group staff of the Washington National Primate Research Center (WaNPRC) for their aid with the RM study. This work was supported by the resources and staff at the University of Minnesota Genomics Center. We thank Dr. Sheri Hild for her support of these studies.

## Author contributions

JAM conceived of the study, oversaw the planning of the project, performed experiments, interpreted the data, and wrote the paper. ATG assisted with sample collection from the RM study, analyzed and interpreted the 16s rRNA sequencing data, and wrote the paper. CAB assisted with sample collection from the RM study and performed the DNA extractions and 16s rRNA sequencing for both the RM and CHMI studies. KH coordinated the collection of fecal samples from the CHMI study. EC provided support for sample processing from the RM study. SW, NI, CYA, and WDG provided research support for the RM study. KAG provided veterinary support for the RM study. KDP provided the initial *P. fragile* stock. MG and NRK assisted with data interpretation and paper preparation. JGK led and provided samples from the KAF156 CHMI study, assisted with data interpretation and paper preparation.

---

**Fig. S1.**
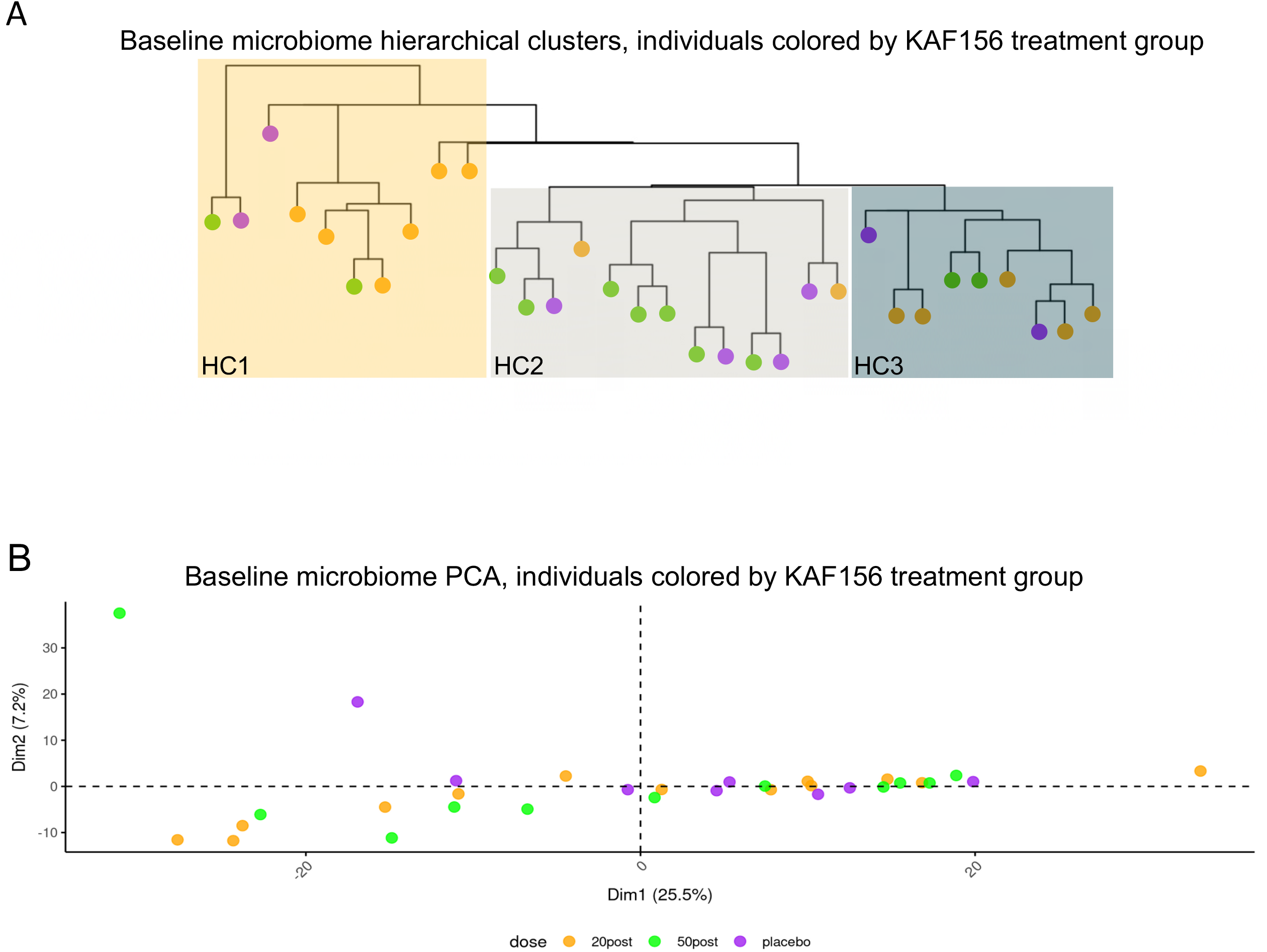
Post-CHMI treatment with KAF156 was not associated with microbiome composition. (A) Divisive hierarchical clustering to identify major clusters in baseline microbiota samples. Major clusters were defined as human cluster 1 (HC1; gold), human cluster 2 (HC2; gray) and human cluster 3 (HC3; dark cyan). Branch tips indicate the KAF156 treatment arm, including placebo (purple), 20mg (orange), or 50mg (green) of KAF156 treatment post-CHMI. (B) Principal component analysis (PCA) of CLR-transformed count data prior to challenge. Colors indicate the KAF156 treatment arm, including placebo (purple), 20mg (orange), or 50mg (green) of KAF156 treatment post-CHMI.

**Fig. S2.**
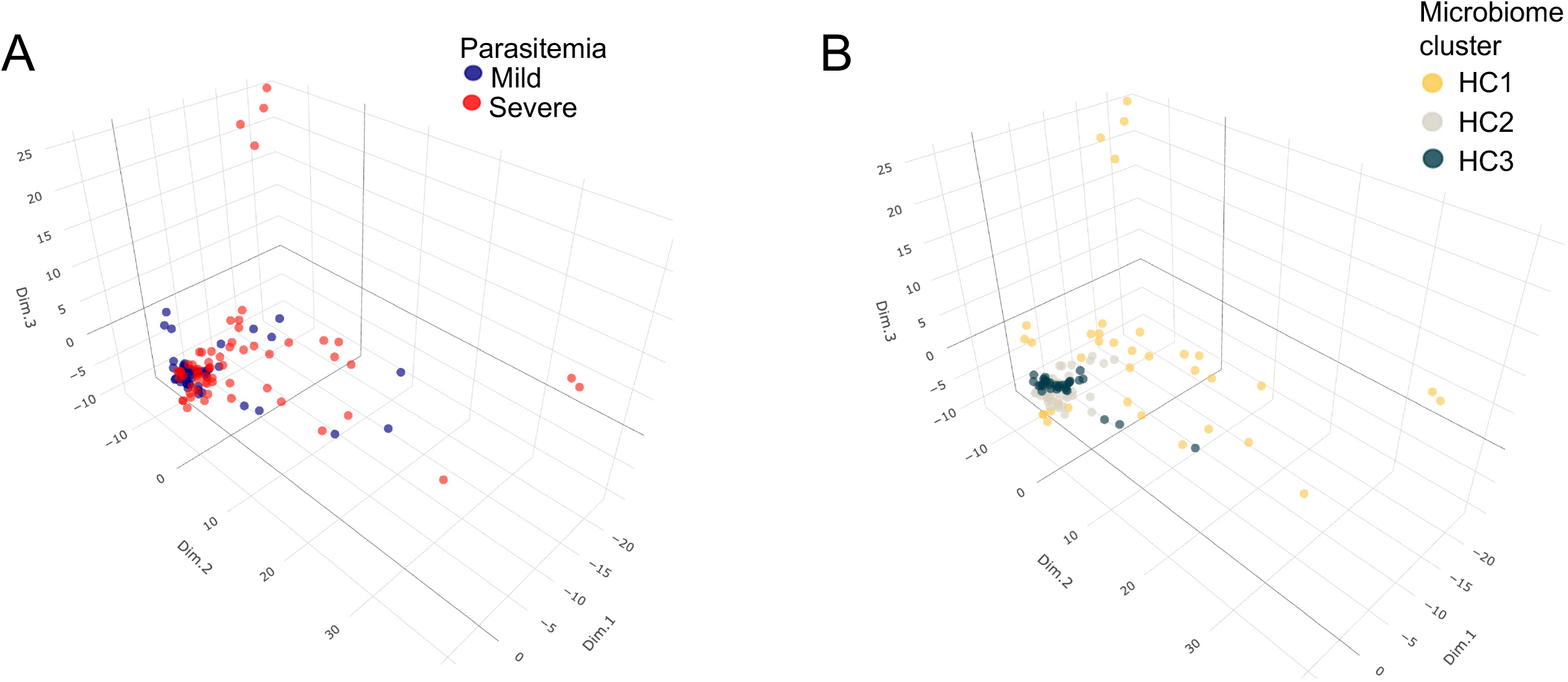
High diversity human GI microbiota linked to likelihood of developing severe *P. falciparum* parasitemia. (A) Three-dimensional principal component analysis (PCA) of CLR-transformed count data, with colors identifying samples associated with parasite levels. Blue points indicate mild parasitemia, while red points indicate severe parasitemia. (B) Replicate PCA shown in (A), with colors indicating hierarchically determined microbiota clusters. HC1 is shown in gold, HC2 is shown in gray and HC3 is shown in dark cyan.

**Fig. S3.**
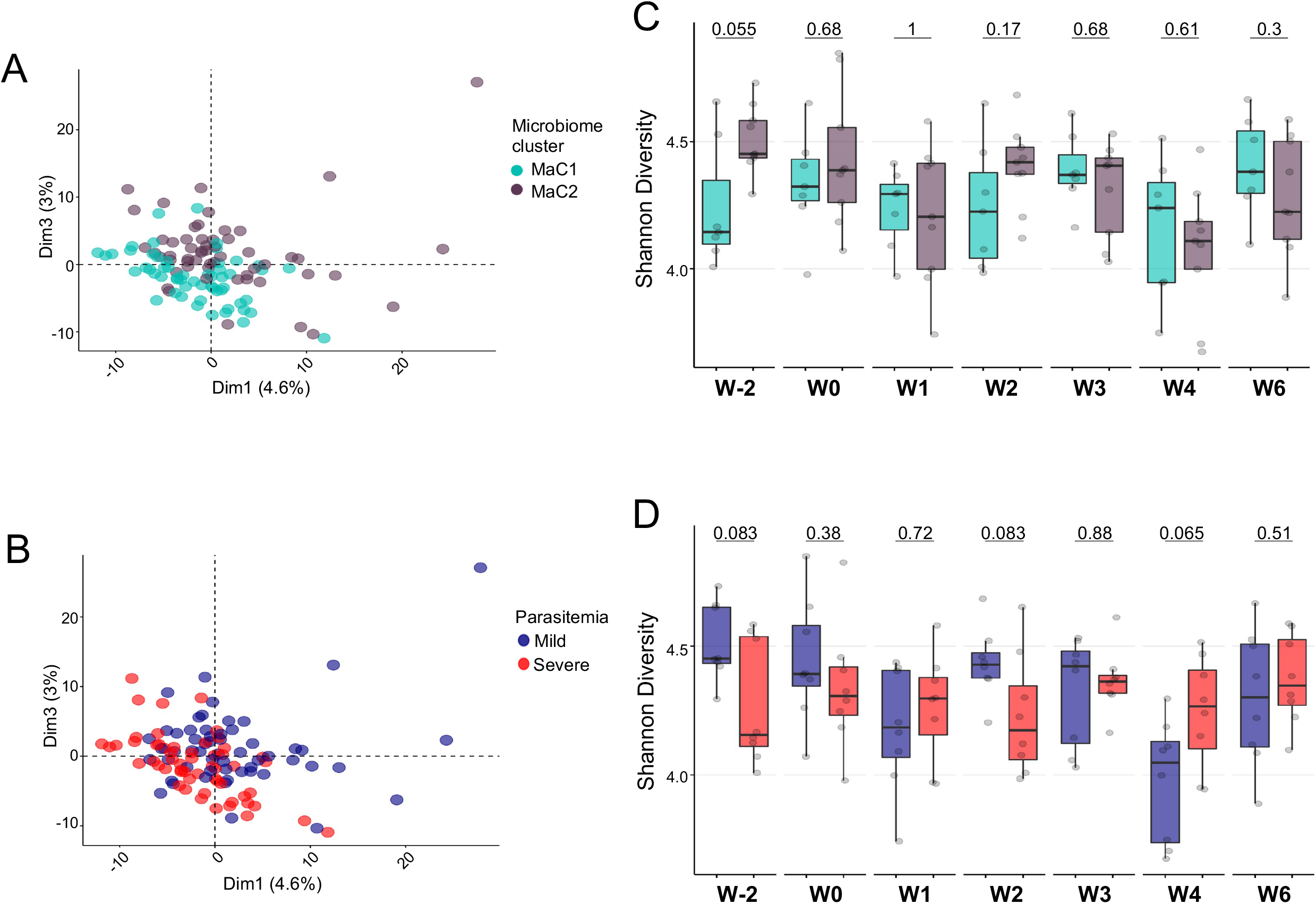
High diversity macaque GI microbiota linked to likelihood of developing severe *P. falciparum* parasitemia. (A) Principal component analysis (PCA) of CLR-transformed count data. Colors indicate hierarchically determined microbiota clusters, with MaC1 shown in teal and MaC2 shown in purple. (B) Replicate PCA shown in (A), with colors identifying samples associated with parasite levels. Blue points indicate mild parasitemia, while red points indicate severe parasitemia. (C) Shannon diversity scores for RM samples at all time points, grouped by microbiota cluster. (D) Shannon diversity scores for RM samples at all time points, grouped by parasitemia severity.

**Fig. S4.**
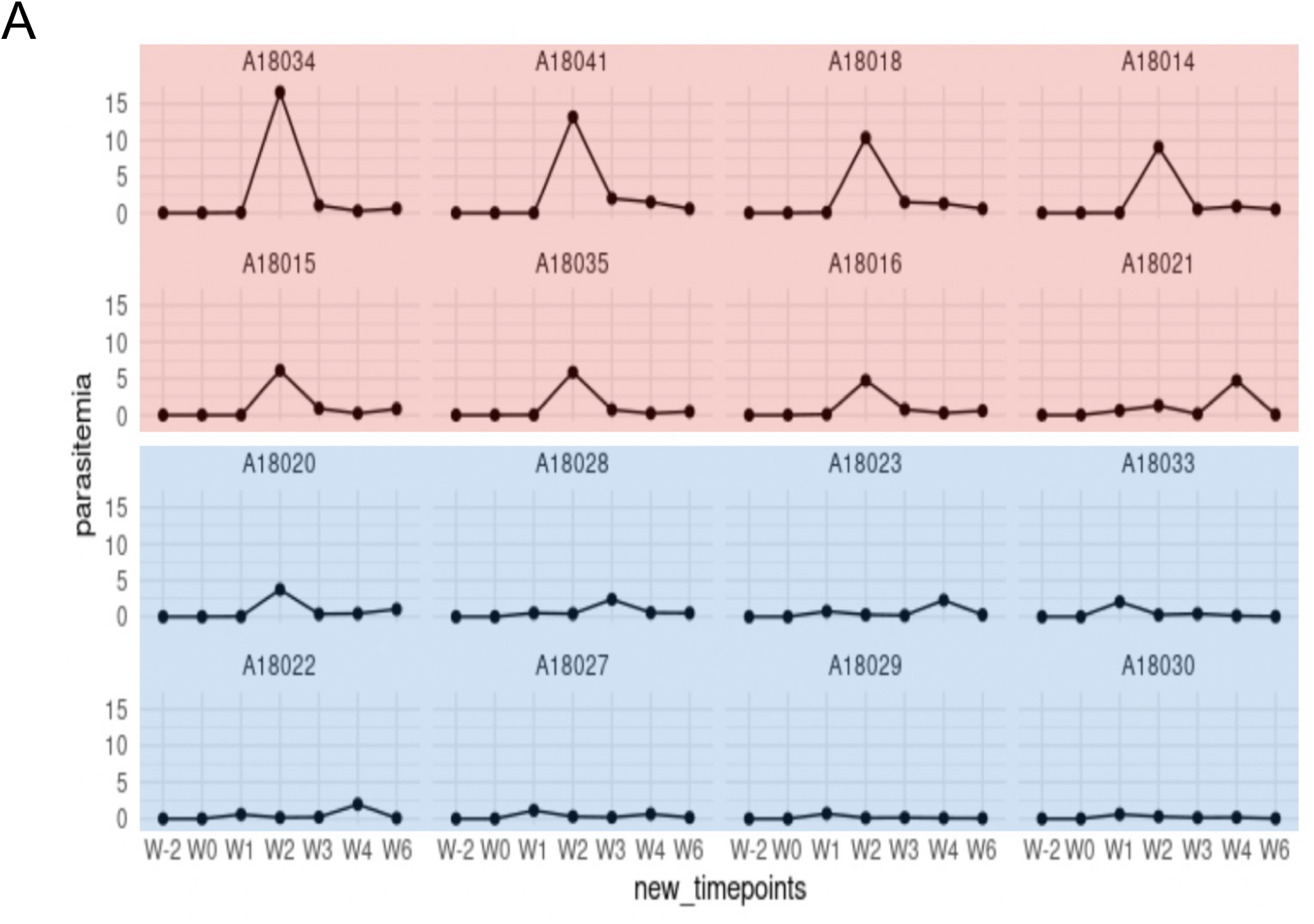
Peak *P. fragile* parasite burdens varied across RMs. (A) *P. fragile* parasite detection in macaques over the study time course. Colors identify samples associated with parasite levels. Blue background indicates RMs with mild parasitemia, while red points indicate RMs that experienced severe parasitemia.

## References

1 WHO. World malaria report 2021. Vol. Licence: CC BY-NC-SA 3.0 IGO (World Health Organization, Geneva, Switzerland, 2021).

2 Sowunmi, A., Ogundahunsi, O. A., Falade, C. O., Gbotosho, G. O. & Oduola, A. M. Gastrointestinal manifestations of acute falciparum malaria in children. Acta tropica 74, 73–76 (2000).

3 Seydel, K. B., Milner, D. A., Jr., Kamiza, S. B., Molyneux, M. E. & Taylor, T. E. The distribution and intensity of parasite sequestration in comatose Malawian children. The Journal of infectious diseases 194, 208–205 (2006). https://doi.org:10.1086/505078

4 Pongponratn, E., Riganti, M., Punpoowong, B. & Aikawa, M. Microvascular sequestration of parasitized erythrocytes in human falciparum malaria: a pathological study. The American journal of tropical medicine and hygiene 44, 168–175 (1991).

5 Mooney, J. P. et al. Inflammation-associated alterations to the intestinal microbiota reduce colonization resistance against non-typhoidal Salmonella during concurrent malaria parasite infection. Scientific reports 5, 14603 (2015). https://doi.org:10.1038/srep14603

6 Milner, D. A., Jr. et al. Quantitative Assessment of Multiorgan Sequestration of Parasites in Fatal Pediatric Cerebral Malaria. The Journal of infectious diseases 212, 1317–1321 (2015). https://doi.org:10.1093/infdis/jiv205

7 Olsson, R. A. & Johnston, E. H. Histopathologic changes and small-bowel absorption in falciparum malaria. The American journal of tropical medicine and hygiene 18, 355–359 (1969).

8 Wilairatana, P., Meddings, J. B., Ho, M., Vannaphan, S. & Looareesuwan, S. Increased gastrointestinal permeability in patients with Plasmodium falciparum malaria. Clinical infectious diseases: an official publication of the Infectious Diseases Society of America 24, 430–435 (1997).

9 Karney, W. W. & Tong, M. J. Malabsorption in Plasmodium falciparum malaria. The American journal of tropical medicine and hygiene 21, 1–5 (1972).

10 Molyneux, M. E. et al. Reduced hepatic blood flow and intestinal malabsorption in severe falciparum malaria. The American journal of tropical medicine and hygiene 40, 470–476 (1989).

11 Church, J. A., Nyamako, L., Olupot-Olupot, P., Maitland, K. & Urban, B. C. Increased adhesion of Plasmodium falciparum infected erythrocytes to ICAM-1 in children with acute intestinal injury. Malaria Journal 15, 54 (2016). https://doi.org:10.1186/s12936-016-1110-3

12 Chau, J. Y. et al. Malaria-associated L-arginine deficiency induces mast cell-associated disruption to intestinal barrier defenses against nontyphoidal Salmonella bacteremia. Infection and immunity 81, 3515–3526 (2013). https://doi.org:10.1128/iai.00380-13

13 Olupot-Olupot, P. et al. Endotoxaemia is common in children with Plasmodium falciparum malaria. BMC infectious diseases 13, 117 (2013). https://doi.org:10.1186/1471-2334-13-117

14 Berkley, J. A. et al. HIV infection, malnutrition, and invasive bacterial infection among children with severe malaria. Clinical infectious diseases: an official publication of the Infectious Diseases Society of America 49, 336–343 (2009). https://doi.org:10.1086/600299

15 Bronzan, R. N. et al. Bacteremia in Malawian children with severe malaria: prevalence, etiology, HIV coinfection, and outcome. The Journal of infectious diseases 195, 895–904 (2007). https://doi.org:10.1086/511437

16 Alamer, E. et al. Dissemination of non-typhoidal Salmonella during Plasmodium chabaudi infection affects anti-malarial immunity. Parasitol Res 118, 2277–2285 (2019). https://doi.org:10.1007/s00436-019-06349-z

17 Potts, R. A. et al. Mast cells and histamine alter intestinal permeability during malaria parasite infection. Immunobiology 221, 468–474 (2016). https://doi.org:10.1016/j.imbio.2015.11.003

18 Roth, A. N., Grau, K. R. & Karst, S. M. Diverse Mechanisms Underlie Enhancement of Enteric Viruses by the Mammalian Intestinal Microbiota. Viruses 11, 760 (2019).

19 Mandal, R. K. et al. Dynamic modulation of spleen germinal center reactions by gut bacteria during Plasmodium infection. Cell Rep 35, 109094 (2021). https://doi.org:10.1016/j.celrep.2021.109094

20 Yooseph, S. et al. Stool microbiota composition is associated with the prospective risk of Plasmodium falciparum infection. BMC genomics 16, 631 (2015). https://doi.org:10.1186/s12864-015-1819-3

21 Villarino, N. F. et al. Composition of the gut microbiota modulates the severity of malaria. Proceedings of the National Academy of Sciences of the United States of America 113, 22352240 (2016). https://doi.org:10.1073/pnas.1504887113

22 Taniguchi, T. et al. Plasmodium berghei ANKA causes intestinal malaria associated with dysbiosis. Scientific reports 5, 15699 (2015). https://doi.org:10.1038/srep15699

23 Waide, M. L. et al. Gut Microbiota Composition Modulates the Magnitude and Quality of Germinal Centers during Plasmodium Infections. Cell Rep 33, 108503 (2020). https://doi.org:10.1016/j.celrep.2020.108503

24 Kublin, J. G. et al. Safety, Pharmacokinetics, and Causal Prophylactic Efficacy of KAF156 in a Plasmodium falciparum Human Infection Study. Clinical infectious diseases: an official publication of the Infectious Diseases Society of America 73, e2407–e2414 (2021). https://doi.org:10.1093/cid/ciaa952

25 Murphy, S. C., Daza, G., Chang, M. & Coombs, R. Laser cutting eliminates nucleic acid cross-contamination in dried-blood-spot processing. Journal of clinical microbiology 50, 4128–4130 (2012). https://doi.org:10.1128/jcm.02549-12

26 Seilie, A. M. et al. Beyond Blood Smears: Qualification of Plasmodium 18S rRNA as a Biomarker for Controlled Human Malaria Infections. The American journal of tropical medicine and hygiene 100, 1466–1476 (2019). https://doi.org:10.4269/ajtmh.19-0094

27 Methods in malaria research. 6th Edition edn, (EVIMalaR and MR4/ATCC, 2013).

28 Lim, C. et al. Improved light microscopy counting method for accurately counting Plasmodium parasitemia and reticulocytemia. Am J Hematol 91, 852–855 (2016). https://doi.org:10.1002/ajh.24383

29 Sim, K. et al. Improved detection of bifidobacteria with optimised 16S rRNA-gene based pyrosequencing. PloS one 7, e32543 (2012). https://doi.org:10.1371/journal.pone.0032543

30 Thompson, L. R. et al. A communal catalogue reveals Earth’s multiscale microbial diversity. Nature 551, 457–463 (2017). https://doi.org:10.1038/nature24621

31 Apprill, A., McNally, S., Parsons, R. & Weber, L. Minor revision to V4 region SSU rRNA 806R gene primer greatly increases detection of SAR11 bacterioplankton. Aquatic Microbial Ecology 75, 129–137 (2015).

32 Caporaso, J. G. et al. Ultra-high-throughput microbial community analysis on the Illumina HiSeq and MiSeq platforms. The ISME journal 6, 1621–1624 (2012). https://doi.org:10.1038/ismej.2012.8

33 Gohl, D. M. et al. Systematic improvement of amplicon marker gene methods for increased accuracy in microbiome studies. Nature Biotechnology 34, 942–949 (2016). https://doi.org:10.1038/nbt.3601

34 Dissanaike, A., Nelson, P. & Garnham, P. Two New Malaria Parasites, Plasmodium cynomolgi ceylonesis subsp. nov and Plasmodium fragile sp. nov., from Monkeys in Ceylon. Ceylon Journal of Medical Research, 717–721 (1965).

35 Collins, W. E. et al. Studies on sporozoite-induced and chronic infections with Plasmodium fragile in Macaca mulatta and New World monkeys. J Parasitol 92, 1019–1026 (2006). https://doi.org:10.1645/ge-848r.1

36 Collins, W. E., Chin, W. & Skinner, J. C. Plasmodium fragile and Macaca mulatta monkeys as a model system for the study of malaria vaccines. The American journal of tropical medicine and hygiene 28, 948–954 (1979). https://doi.org:10.4269/ajtmh.1979.28.948

37 Abt, M. C. et al. Commensal bacteria calibrate the activation threshold of innate antiviral immunity. Immunity 37, 158–170 (2012). https://doi.org:10.1016/j.immuni.2012.04.011

38 Brown, R. L., Sequeira, R. P. & Clarke, T. B. The microbiota protects against respiratory infection via GM-CSF signaling. Nature communications 8, 1512 (2017). https://doi.org:10.1038/s41467-017-01803-x

39 Ganal, S. C. et al. Priming of natural killer cells by nonmucosal mononuclear phagocytes requires instructive signals from commensal microbiota. Immunity 37, 171–186 (2012). https://doi.org:10.1016/j.immuni.2012.05.020

40 Hensley-McBain, T. et al. Increased mucosal neutrophil survival is associated with altered microbiota in HIV infection. PLoS pathogens 15, e1007672 (2019). https://doi.org:10.1371/journal.ppat.1007672

41 Rooks, M. G. & Garrett, W. S. Gut microbiota, metabolites and host immunity. Nat Rev Immunol 16, 341–352 (2016). https://doi.org:10.1038/nri.2016.42

42 Yang, Y. et al. Within-host evolution of a gut pathobiont facilitates liver translocation. Nature 607, 563–570 (2022). https://doi.org:10.1038/s41586-022-04949-x

43 Mandal, R. K. et al. Longitudinal Analysis of Infant Stool Bacteria Communities Before and After Acute Febrile Malaria and Artemether-Lumefantrine Treatment. The Journal of infectious diseases 220, 687–698 (2019). https://doi.org:10.1093/infdis/jiy740

44 Guan, W. et al. Observation of the Gut Microbiota Profile in BALB/c Mice Induced by Plasmodium yoelii 17XL Infection. Frontiers in microbiology 13, 858897 (2022). https://doi.org:10.3389/fmicb.2022.858897

45 Guan, W. et al. Observation of the Gut Microbiota Profile in C57BL/6 Mice Induced by Plasmodium berghei ANKA Infection. Frontiers in cellular and infection microbiology 11, 680383 (2021). https://doi.org:10.3389/fcimb.2021.680383

46 Fan, Z. G. et al. Gut Microbiota Reconstruction Following Host Infection with Blood-stage Plasmodium berghei ANKA Strain in a Murine Model. Curr Med Sci 39, 883–889 (2019). https://doi.org:10.1007/s11596-019-2119-y

47 Zmora, N., Suez, J. & Elinav, E. You are what you eat: diet, health and the gut microbiota. Nature reviews. Gastroenterology & hepatology 16, 35–56 (2019). https://doi.org:10.1038/s41575-018-0061-2

